# Linear DNA amplicons as a novel cancer vaccine strategy

**DOI:** 10.1101/2022.02.09.479777

**Authors:** Antonella Conforti, Erika Salvatori, Lucia Lione, Mirco Compagnone, Eleonora Pinto, Clay Shorrock, James A. Hayward, Yuhua Sun, Ben Minghwa Liang, Fabio Palombo, Brian Viscount, Luigi Aurisicchio

**Affiliations:** Takis Biotech, Via Castel Romano 100, 00128 Rome, Italy; Evvivax Biotech, Via Castel Romano 100, 00128 Rome, Italy; Neomatrix Biotech, Via Castel Romano 100, 00128 Rome, Italy; Applied DNA Sciences, 50 Health Sciences Drive, Stony Brook, NY 11790

**Keywords:** DNA vaccine, PCR, amplicons, cancer immunotherapy

## Abstract

DNA-based vaccines represent a simple, safe and promising strategy for harnessing the immune system to fight infectious diseases as well as various forms of cancer and thus are considered an important tool in the cancer immunotherapy toolbox. Nonetheless, the manufacture of plasmid DNA vaccines has several drawbacks, including long lead times and the need to remove impurities from bacterial cultures. Here we report the development of polymerase chain reaction (PCR)-produced amplicon expression vectors as DNA vaccines and their *in vivo* application to elicit antigen-specific immune responses in animal cancer models. Amplicons encoding tumor-associated-antigens, such as telomerase reverse transcriptase or neoantigens expressed by murine tumor cell lines were able to elicit antigen-specific immune responses and proved to significantly impact tumor growth when administered in combination with immune-checkpoint inhibitors (ICIs). These results strongly support the further exploration of the use of PCR-based amplicons as an innovative immunotherapeutic approach to cancer treatment.

## Introduction

Cancer is the leading cause of mortality worldwide. Conventional therapies such as surgery, radiation and chemotherapy, are considered aggressive and can negatively impact the patient without offering lifelong protection, immunotherapy has in recent years emerged as a promising, more tolerable and more durable form of cancer treatment. Among the newest immunotherapeutic approaches to cancer treatment, DNA-based cancer vaccines are important strategies in the cancer immunotherapy toolbox.^1^ DNA vaccines for cancer immunotherapy are designed to deliver one or several genes encoding tumor-associated antigens or other immunogenic polypeptides to modulate immune responses, thereby eliciting or strengthening immune responses against tumor antigens that play a central role in tumor initiation, progression and metastasis, or arise as a result of mutational burden, e.g. neoantigens.^2,3,4,5^ DNA cancer vaccines can induce both innate immunity and adaptive immune responses which can suppress tumor growth and, in some cases, achieve total tumor rejection or shrinkage.^6,7,8,9^ To date, DNA based vaccines have involved the inoculation of a subject with plasmid DNA and/or plasmid-derived DNA. Bacterial plasmids are episomal circular DNA constructs propagated in bacteria via a fermentation process. However, the manufacture and use of DNA vaccines via plasmids has several drawbacks, including the presence of antibiotic resistance genes, the presence of unwanted additional DNA sequences in the form of the plasmid backbone, and the need to remove impurities from bacterial cultures, such as bioburden and endotoxins, long lead times, inefficient uptake of the large plasmid DNA molecules to the cellular nucleus and challenges of integrating plasmid production into automated cGMP (current Good Manufacturing Practices) workflows. Thus, there is an unmet need for a new DNA vaccination platform not including bacterial plasmids or plasmid-derived DNA. In this study, we report the development of an innovative DNA vaccine based on a DNA amplicon produced by the enzymatic amplification process of an expression vector via the polymerase chain reaction (PCR). In order to improve the weak immunogenicity induced by naked DNA injection, we exploited electroporation (EP) as a powerful delivery technology, considering its safety, low cost, efficacy and ease of application. This procedure consists of exposing a cell or a tissue to an external electric field that increases cell membrane permeability to molecules that otherwise would cross the plasma membrane with low efficiency; for this reason, it has been widely used in different biomedical applications.^10,11,12^ Comparison between DNA injection alone and DNA-EP has demonstrated an increase in both cellular and humoral response after electric fields were applied. It has been demonstrated that the addition of electroporation provides a 10-100-fold augmentation of immune response and defense against pathogens in humans and numerous animal models of diseases such as HIV/SIV, malaria, hepatitis B and C, human papilloma virus (HPV), anthrax and influenza.^13^ Furthermore, EP may enhance immune responses through increased protein expression, secretion of inflammatory chemokines and cytokines, and recruitment of monocytes and antigen-presenting cells at the site of electroporation.^14,15^ In addition, *in vivo* electroporation causes transient and reversible cell damage also resulting in local inflammation and release of cytokines which may act as a danger signal, consequently enhancing vaccine potency.^16,17^ Here, we demonstrate gene expression mediated by DNA amplicons both *in vitro* and *in vivo* and show the immunogenicity of a DNA amplicon encoding for a tumor associated antigen (i.e. Telomerase Reverse Transcriptase - TERT) in wild-type mice as compared to plasmid DNA-EP. Furthermore, we show that DNA amplicons encoding neoantigens and delivered by EP induce therapeutic effects in tumor-bearing mice, when combined with Immune Checkpoint Inhibitors (ICIs), in comparison to the combination of plasmid DNA-EP and ICIs.

## Materials and Methods

### Plasmid DNA constructs

All constructs were completely synthesized and optimized for codon usage. Synthesis and codon optimization analysis of plasmid DNA vectors were performed at Genscript (China).

Luciferase gene, cloned into the linearized pcDNA3-Hygro vector by enzymatic restriction, was used to assess gene expression both *in vitro* (by transfecting cells) and *in vivo* (by mice immunization).

conTRT vaccine consisted of a DNA plasmid encoding a consensus sequence between canine and human telomerase. In addition, this catalytically inactive telomerase protein has been fused to a TPA (human tissue plasminogen activator) leader sequence at N-term and fused itself to the Profilin-like sequence of *Toxoplasma gondii* (PFTG) at the C-terminus (TPA-conTRT-PFTG_opt_). This construct was finally cloned into the linearized pTK1A-TPA vector by enzymatic restriction. Neoantigen-based vaccines are DNA plasmids encoding for neoepitopes selected from a murine colon adenocarcinoma cell line (i.e., MC38 cell line, ATCC, USA). Once identified by RNA sequencing, genes encoding for selected neoepitopes were cloned in pTK1A vector, as previously described.^7,18^

### DNA Amplicon Constructs

PCR primers were designed to amplify the expression cassette contained in the plasmid sequences for the Luciferase, conTRT vaccine and Neoantigen-based vaccine (M8) constructs, so creating dsDNA amplicons consisting only of the expression cassette sequence (namely, DNA amplicon expression vector). For the phosphothiate-modified amplicons, a sulfur atom substitutes the non-bridging oxygen in the phosphate backbone of the oligonucleotide. In addition, the three (3’) terminal bases of both the forward and reverse primers were modified to increase DNA amplicon stability.^19^ DNA amplifications via PCR for the Luciferase constructs were performed using GeneAmp 2700 PCR system (Applied Biosystem, USA) using either BIOLASE^TM^ polymerase (7×10^-5^ approximate error/bp) or MyFi^TM^ polymerase (7×10^-5^ approximate error/bp) both from Bioline (Meridian Bioscience, USA). The PCR mixture consisted of 1X buffer with 1mM dNTP, 0.5μM of forward and reverse primers, polymerase 5u/100ul, and plasmid template at 40ng/mL. The PCR reactions were subjected to initial denaturation of 97°C for 20 seconds, and subsequent 30 cycles of 95°C for 10 seconds and 72°C for 5 minutes followed by final extension at 72°C for 3 minutes. Four different versions of Luciferases DNA amplicon expression vectors were produced using either regular primers or phosphothiate-modified primers using either the Biolase or MyFi polymerase, resulting in 2878 bp DNA amplicon expression vectors. DNA amplifications for the conTRT amplicons were performed similarly except Ranger DNA polymerase (1×10^-6^ approximate error/bp) from Bioline was utilized, resulting in 6024 bp DNA amplicon expression vectors. DNA amplifications for the M8 amplicon were also performed similarly except Q5^TM^ polymerase (5.3×10^-7^ approximate error/bp; New England BioLabs, USA) was utilized, resulting in 3037 bp DNA amplicon expression vectors.

### DNA purification and qualification

Each dsDNA amplicon expression vector produced via the PCR amplification was purified on an Akta Pure 150 FPLC instrument (GE Healthcare, USA). After the entire lot was loaded onto the GE ReadyToProcess Adsorber Q anion exchange column and washed, a linear gradient of up to 1.5 M NaCl in 1x PBS was used to elute DNA via 0.5 mL fractions. These fractions were analyzed with DNA 7500 assay on 2100 Bioanalyzer (Agilent, USA). Fractions containing the DNA amplicon expression vectors of interest were identified, pooled, and concentrated using ethanol precipitation and refrigerated centrifugation. After rinsing and drying, each DNA amplicon expression vector was resuspended to the desired concentration in 1x PBS, and sterile filtered. Further analytical characterization of each quantity of DNA amplicon expression vector was performed using NanoDrop (Thermo Fisher, USA), 2100 Bioanalyzer, and Alliance HPLC System (Waters, USA).

Based on the above workflows, the following DNA expression vectors (DNA amplicons) were produced: as for Luciferase, #349 (PS-modified; MyFi Polymerase), #351 (PS-modified; Biolase Polymerase), #355 (MyFi Polymerase), #358 (Biolase Polymerase); as for conTRT, #359 (PS-modified; Ranger Polymerase), #360 (PS-modified; Biolase Polymerase), #363 (Ranger Polymerase), #365 (Biolase Polymerase); as for M8, #12290 (Q5 Polymerase).

### Vaccination and murine models

For *in vivo* expression and immunogenicity assessment, 6-8 weeks old BALB/c female mice (Envigo, USA) were injected intramuscularly (i.m.), particularly in the quadriceps, with either DNA plasmid or amplicon (dose ranging from 5 μg to 50 μg) and electrically stimulated as previously described.^20,21^ The DNA was formulated in Phosphate Buffered Saline (PBS) at a concentration of 0.1-1 mg/ml. DNA-EP was performed by means of a Cliniporator Device EPS01 and using N-10-4B electrodes (IGEA, Italy) with the following electrical conditions in Electro-Gene-Transfer (EGT) modality: 8 pulses 20 msec each at 110V, 8Hz, 120msec interval. As for the evaluation of the antitumoral efficacy of plasmid and amplicon DNA in cancer models, tumor challenge was performed by injecting 3×10^5^ MC38 cells subcutaneously in the right flank of 6-8 weeks old female C57Bl/6 mice (Envigo). Twice a week, mice were examined for tumor growth using a caliper. Tumor volumes were calculated using the formula:

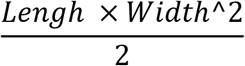

Mice were housed according to national legislation and kept in standard conditions according to Evvivax ethical committee approval. All the *in vivo* experimental procedures were approved by the local animal ethics council.

### Peptides

Lyophilized peptides covering telomerase reverse transcriptase protein or neoepitopes were purchased from JPT (Germany) and resuspended in DMSO at 40 mg/ml. As for telomerase, pools of peptides of 15 aa overlapping by 11 residues were assembled in four pools (A, B, C, D). Peptides and pools were stored at −80°C.

### Luciferase assay

*In vitro* luciferase assay was performed by cell transfection. Briefly, Hela cells were plated at 1×10^5^/well in 96well plate and then transfected with equimolar concentrations of DNA plasmid or DNA amplicon (1 and 0.3 μg). After 24 hours, luciferase expression signal was measured by Imaging at an IVIS 200 imaging device (Perkin Elmer, USA).

In order to assess luciferase expression *in vivo,* female BALB/c mice (5 mice/group) were anesthetized using 97% oxygen and 3% isoflurane (Isoba, UK) then injected by EP in quadriceps muscle with a DNA plasmid encoding Luciferase (pcDNA3-Hygro-Luc at 1 μg/mouse) or equimolar concentrations of DNA amplicon. Imaging was performed under gas anesthesia at IVIS 200 imaging system at 24h, 48h and one week after injection, 8 min after injecting subcutaneously a luciferin substrate solution (15mg/ml, Perkin Elmer) at 10μl/g of body weight.

### Flow Cytometry analysis

In experiments performed with vaccinated mice, the intracellular staining was performed according to the procedure described in Giannetti et al.^22^ Briefly, PBMC or splenocytes were treated with ACK Lysing buffer (Life Technologies, USA) for red blood cell lysis and resuspended in 0.6ml RPMI, 10% FCS and incubated with the indicated pool of peptides (5 μg/ml final concentration of each pool) and brefeldin A (1 μg/ml; BD Pharmingen, USA) at 37°C for 12-16 hours. Cells were then washed and stained with surface antibodies. After washing, cells were fixed, permeabilized and incubated with anti-IFNγ (XMG1.2) and anti-TNFα (MP6-XT22; all from eBioscience, USA), fixed with 1% formaldehyde in PBS, acquired by means of a CytoFLEX LX flow cytometer (Beckman Coulter, USA) and analysis was performed using CytExpert software (Beckman Coulter). DMSO and PMA/IONO (Sigma, Italy) at 10μg/ml were used as internal negative and positive control of the assay, respectively.

### IFNγ ELISpot

T cell ELISpot for mouse IFNγ was performed as previously described.^23^ Briefly, splenocytes isolated from either conTRT vaccinated BALB/c mice or neoantigen vaccinated mice were stimulated for 20h with telomerase or neoantigen peptide pools (1 μg/ml as final concentration). After developing the assay, spot forming cells (SFCs) were counted using an automated ELISPOT reader (A.EL.VIS ELIspot reader, Germany).

### Statistical analyses

Statistical analyses were performed with GraphPad Prism software version 8 (GraphPad). *n* represents individual mice analyzed per experiments. Error bars indicate the standard error of the mean (SEM). We used Mann-Whitney U-tests to compare two groups with non-normally distributed continuous variables. Significance is indicated as follows: *p<0.05; **p<0.01. Comparisons are not statistically significant unless indicated.

## Results

### Manufacturing of DNA Amplicons

A schematic representation showing the assembly of termini phosphorothioate-modified (PS-modified) amplicon expression vectors via a PCR device is shown (Fig. 1A). The DNA sequence of the amplicon expression vector (taken from plasmid DNA expression cassette sequence) includes a promoter, one open reading frame (ORF) and a terminator. It may optionally include a fusion ORF or other secondary ORFs and/or one or more enhancers. Once the sequence is known, a template amplicon expression vector is created via DNA synthesis, without the use of plasmid DNA. Appropriate PS-modified PCR primers for the specific sequence of the template amplicon expression vector are generated via oligonucleotide synthesis. The template amplicon expression vector and the amine-modified primers are introduced to a PCR device whereby the template amplicon expression vector is exponentially amplified via PCR to create amplicon expression vector with PS-modified 3’ and/or 5’ termini. Upon completion of the PCR reaction the amplicon expression vectors are purified and ready for use in the transfection of a target cell for expression of the desired peptide, antigen, polypeptide or protein.

**Figure 1.**
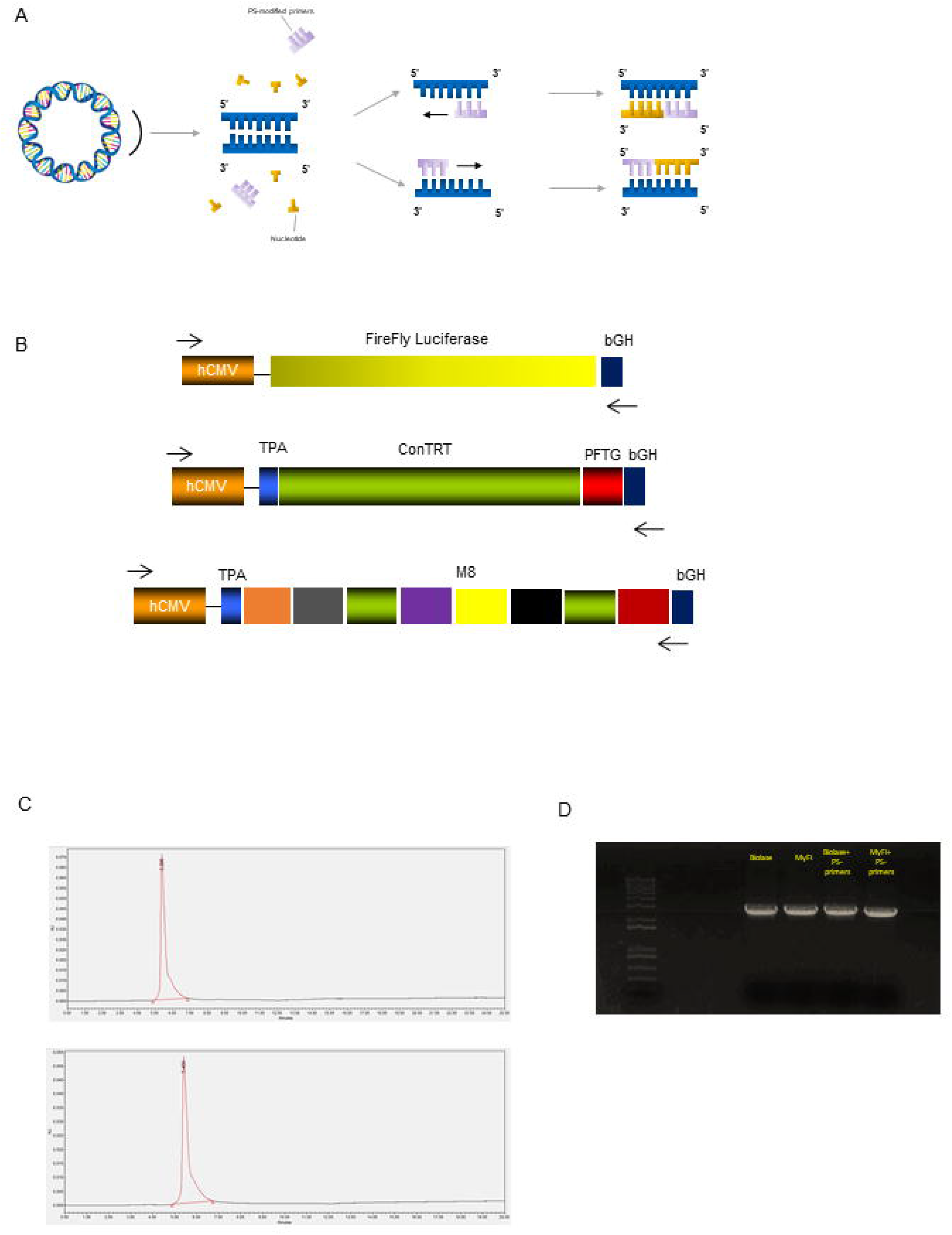
Manufacturing process of DNA amplicons. **(A)** A schematic representation showing the assembly of termini PS-modified amplicon expression vectors via a PCR device is shown. The antigen-encoding sequence, taken from a plasmid DNA, comprises a promotor, one or more ORFs and a terminator. Amplicon expression vector, encoding the selected sequence, is syntethized and then amplified through PS-modified PCR primers in order to produce a plurality of amplicon expression vectors. **(B)** A schematic representation of gene expression cassettes is shown for luciferase, conTRT and M8 constructs. **(C)** HPLC chromatogram of two representative DNA amplicons, one synthetized by means of Biolase^TM^ Taq (top) and one synthetized by means of MyFi^TM^ Taq (bottom). **(D)** Electrophoresis gel of four DNA amplicons, all encoding luciferase gene (synthetized by means of Biolase™ or MyFi^TM^ Taq polymerase, with or w/o PS-modified primers).

As shown in Fig.1B, for the luciferase expression cassette, human CMV promoter drives the expression of firefly luciferase and a bovine growth hormone (bGH) poly A has been used as termination site. conTRT is a codon-optimized, consensus telomerase fused with Profilin-like from *Toxoplasma Gondii* (PFTG). M8 is a polypeptide encoding 8 neoantigens expressed by MC38 murine colon cancer cells.^5,7^ For conTRT and M8 constructs, a human CMV promoter/intron A drives gene expression as well.

Various polymerases, with different error rates, have been used for the PCR-based production of the amplicon expression vectors. BIOLASE^TM^ polymerase (7×10^-5^ approximate error/bp), MyFi^TM^ polymerase (7×10^-5^ approximate error/bp) and RANGER™ polymerase (1×10^-6^ approximate error/bp) have been used to amplify the expression cassette in the presence or absence of PS-modified primers. Figures 1C and 1D show the purity of the amplicons encoding luciferase, measured by HPLC and gel electrophoresis, respectively. These data confirm that the usage of different Taq polymerase, in the presence of PS-modified primers or not, does not affect DNA amplicons purity.

### Assessment of *in vitro* and *in vivo* expression of plasmid and DNA amplicons

In order to compare the expression capacity of DNA amplicons to plasmid DNA expression, the four differentially manufactured PCR products were evaluated by means of *in vitro* cell transfection and *in vivo* injection. For *in vitro* assay, Hela cells were transfected with equimolar concentrations of plasmid DNA (pcDNA3-Hygro-Luc) and four different PCR products (1 and 0.3 μg dose), all encoding luciferase. 24 hours after transfection, luciferase signal was revealed both in plasmid DNA and in DNA amplicon transfected cells, at all tested doses, although at lower extent in latter ones (data not shown). For *in vivo* evaluation of luciferase expression, BALB/c mice were injected i.m. by DNA-EP, with equimolar concentrations of plasmid DNA or DNA amplicons encoding for luciferase (50-10-5 μg/mouse). Luciferase signal was measured by performing optical imaging 24h, 48h and one week post injection (Fig.2A). As shown in Fig.2B, all mice showed luciferase expression in both legs with 50 μg plasmid DNA dose (corresponding to 18.75 μg of DNA amplicon) at each evaluation time point, despite a consistent decrease in mice treated with DNA amplicons at one week post injection. Conversely, mice treated with 50μg of plasmid DNA showed a constantly high luciferase expression. Analysis of counts values confirmed the expression capacity of luciferase-encoding PCR products, although lower than that of plasmid DNA. Interestingly, while luciferase expression by plasmid DNA kept growing from 24h measurement to one week after injection, luciferase signal in mice treated with DNA amplicons reached the highest value at 48h post injection and declined at one week time point, although still being measurable (Fig. 3A). This observation was generally valid also in mice injected with 10 μg of plasmid DNA (corresponding to 3.75 μg of DNA amplicon), although with greater variability (Fig.3B), and in mice injected with 5 μg of DNA amplicon (corresponding to 1.875 μg of PCR product) although with lower counts measured in mice injected with DNA amplicons (Fig.3C). These results show that DNA amplicon expression is stable for at least 48h after being injected in mouse quadriceps, being still detectable even at one week after injection.

**Figure 2.**
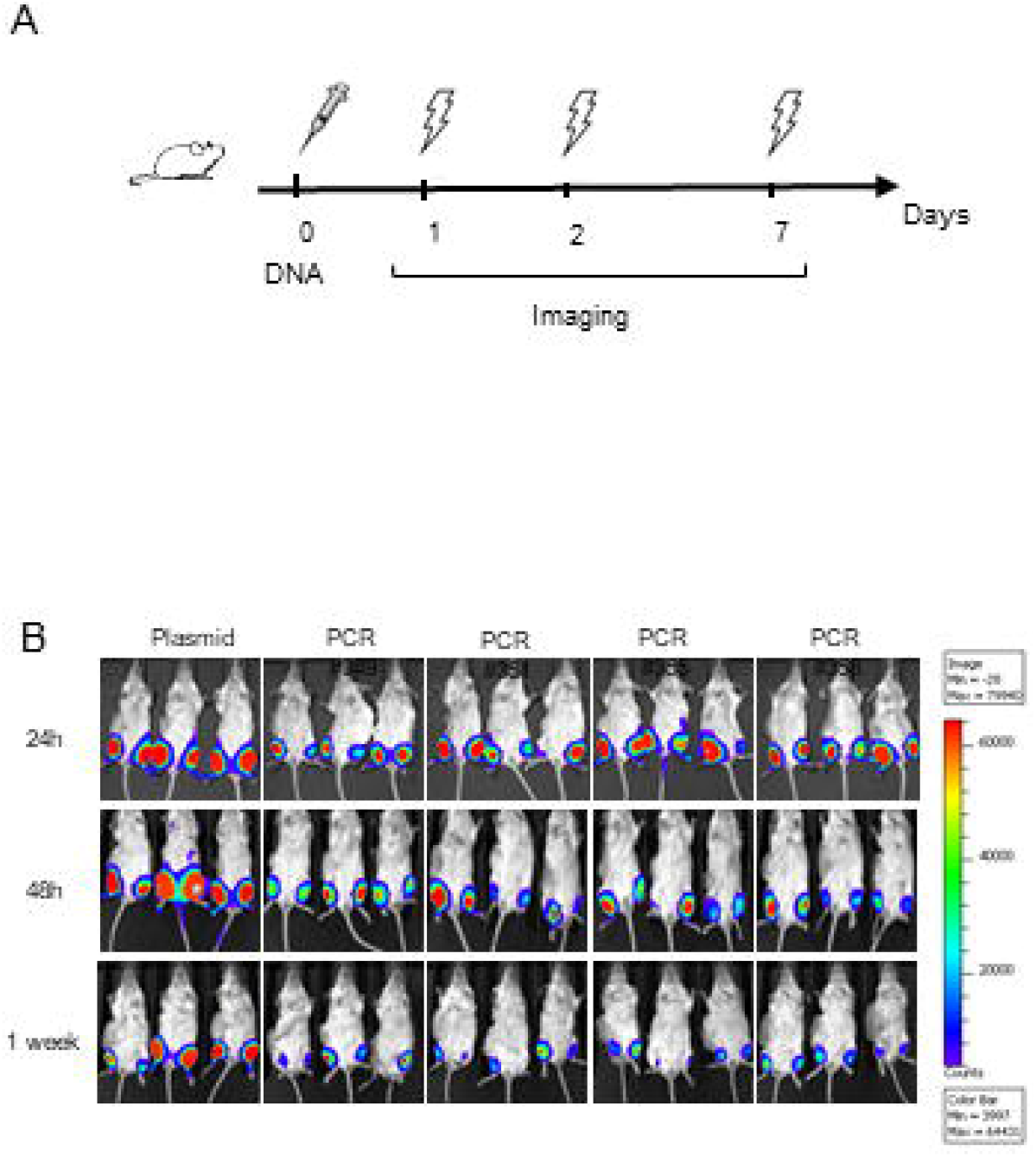
*In vivo* expression of DNA amplicons. **(A)** Schematic representation of the experimental setup. BALB/c mice were injected i.m. by DNA-EP, with equimolar concentrations of plasmid DNA or DNA amplicons encoding for luciferase (50-10-5 μg/mouse). **(B)** After 1-2-7 days post DNA injection, luciferase signal was measured by performing optical imaging at IVIS. Although a decreased expression was detected in DNA amplicons injected mice after 7 days, no difference was measured between DNA amplicons synthetized by means of different Taq polymerase.

**Figure 3.**
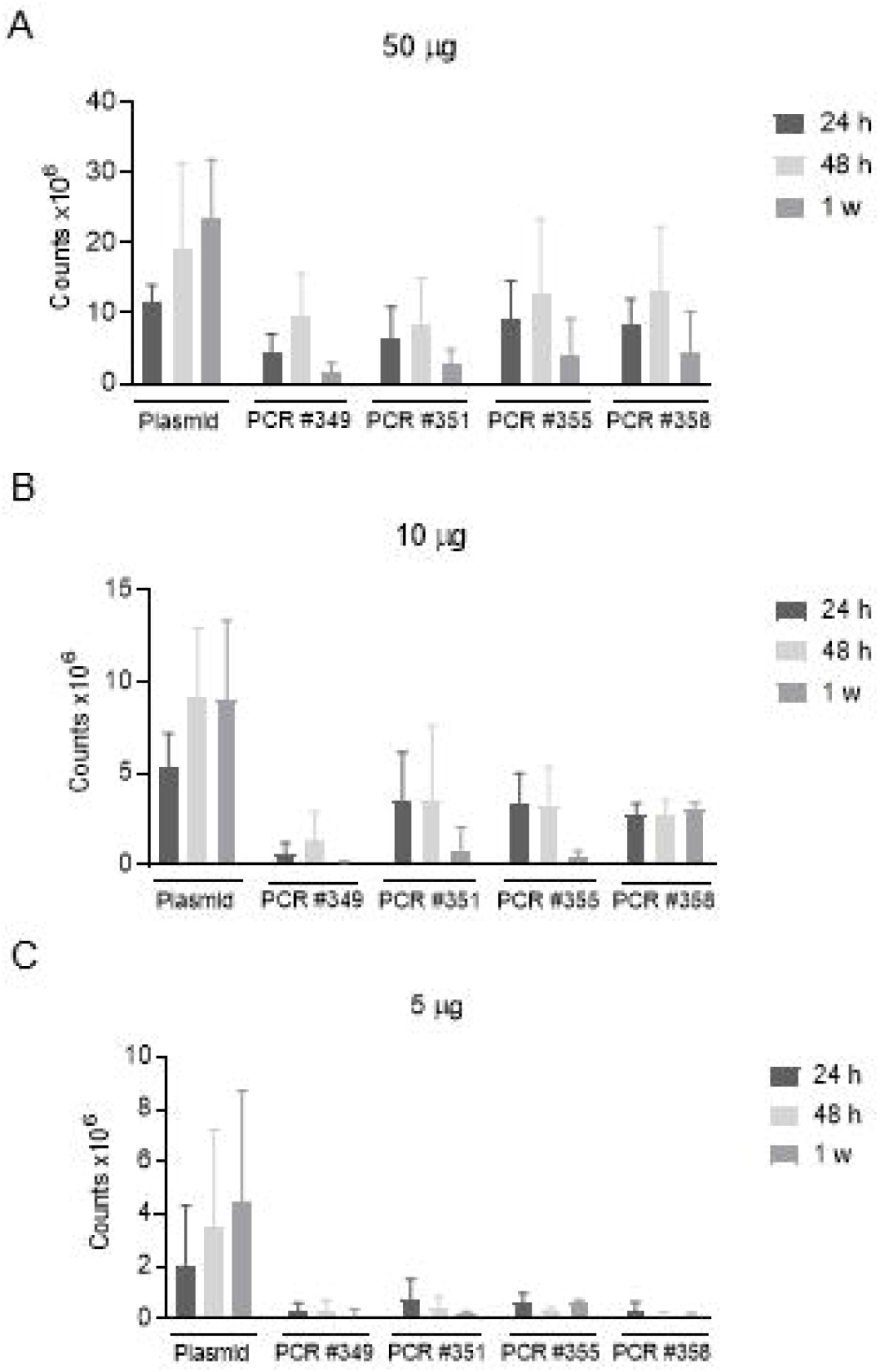
Assessment of *in vivo* DNA amplicon expression. By means of optical imaging at IVIS, in vivo plasmid DNA and amplicon expression was assessed after 24h, 48h and one week post DNA injection at different DNA doses, 50 μg (**A**) – 10 μg (**B)** – 5 μg **(C)**.

### *In vivo* analysis of T cell response elicited by DNA amplicons

Once we demonstrated the expression of DNA amplicons after *in vivo* injection, we sought to evaluate their capacity to elicit an immune response against a tumor associated antigen. To this aim, we vaccinated BALB/c mice i.m. by means of DNA-EP, with three doses (50-10-5 μg/mouse) of a DNA plasmid encoding for conTRT. Likewise, we vaccinated BALB/c mice with equimolar doses (33.75-6.75-3.375 μg/mouse) of a DNA amplicon encoding for the same sequence. All mice were vaccinated following a prime-boost regimen (days 0 and 21, see Fig.4A) and blood analysis was performed at day 28 by intracellular cytokine staining. Cytokine analyses revealed a Th1-skewed immune response, both in plasmid DNA vaccinated mice and DNA amplicon vaccinated mice. For CD8+ T cell response (Fig.4B), there was no significant difference in the magnitude of antigen-specific T cells, at all tested DNA doses, with the only exceptions of mice treated with 10 μg of plasmid DNA and mice treated with 5 μg of amplicon #360. As for CD4+ T cell compartment, no significant differences were revealed between plasmid DNA and DNA amplicons vaccinated mice as well, with the only exception of mice treated with 10μg of plasmid DNA (Fig.4C). These results demonstrate that, although in some cases there is no dose-dependent immunogenicity, DNA plasmid and PCR product induce comparable immune response *in vivo* against a tumor associated antigen.

**Figure 4.**
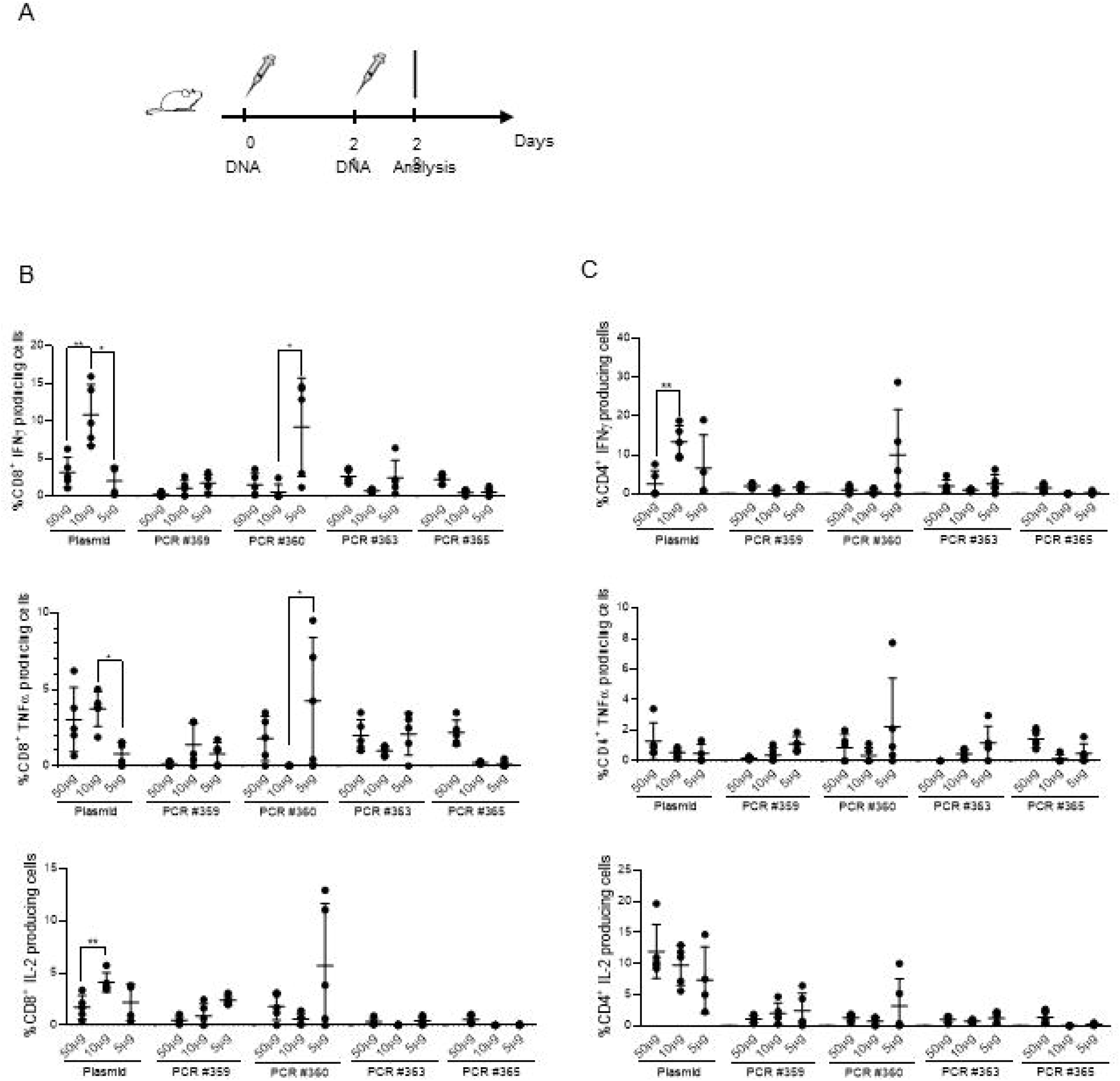
*In vivo* immunogenicity of DNA amplicons. **(A)** Schematic representation of experimental setup. BALB/c mice were injected, by means of i.m. DNA-EP, with three doses (50-10-5 μg/mouse) of a DNA plasmid, and equimolar doses of DNA amplicons, both encoding for conTRT. All mice were vaccinated following a prime-boost regimen (days 0 and 21) and blood analysis was performed at day 28 by intracellular cytokine staining. Assessment of Th1 **(B)** and Th2 **(C)** immune response was done on PBMCs isolated at day 28. Significance was determined using Mann-Whitney test, *p<0,05 **p< 0.01.

### Antitumoral effect of DNA amplicons in murine cancer model

After proving that DNA amplicons are able to elicit antigen-specific immune responses as efficiently as plasmid DNA, we investigated whether this novel immunotherapeutic strategy could exert an antitumoral effect in a cancer murine model. To this aim, we exploited a murine cancer model previously described,^5^ where a neoantigen-based DNA vaccine (namely, M8) was shown to induce tumor regression when administered in combination with ICIs such as αPD1 and αCTLA4, in a therapeutic setting. Briefly, as shown in Fig.5A, after tumor challenge, C57Bl/6 mice received a combination of DNA (plasmid or amplicon, administered at days 2,9 and 16) and ICIs (αCTLA-4 or αPD1, administered at days 3,6 and 9). Neoantigen-specific T cell response, assessed by IFNγ ELISpot performed on splenocytes collected at day 23, was significantly increased over the control mice after cotreatment with plasmid M8 and αCTLA-4 or both ICIs, but not in mice cotreated with plasmid M8 and αPD1, thus suggesting a synergistic effect of αCTLA-4 and plasmid M8 vaccine in eliciting an antitumoral effect in this cancer model. Interestingly, this synergistic effect was confirmed when M8 amplicon was administered in combination with αCTLA-4 (Fig. 5B). Furthermore, the cotreatment of M8 amplicon and both ICIs was significantly more efficient in eliciting a neoantigen-specific T cell response than the cotreatment of plasmid M8 and both ICIs. This observation was confirmed by the assessment of tumor growth (Fig.5C), measured until day of sacrifice (day 23). As observed in mice cotreated with plasmid M8 and ICIs, also the cotreatment with M8 amplicon and ICIs was equally effective in slowing down tumor progression over control mice. These data further support the potential use of linear DNA as alternative to plasmid DNA in developing immunotherapeutic strategies against cancer disease.

**Figure 5.**
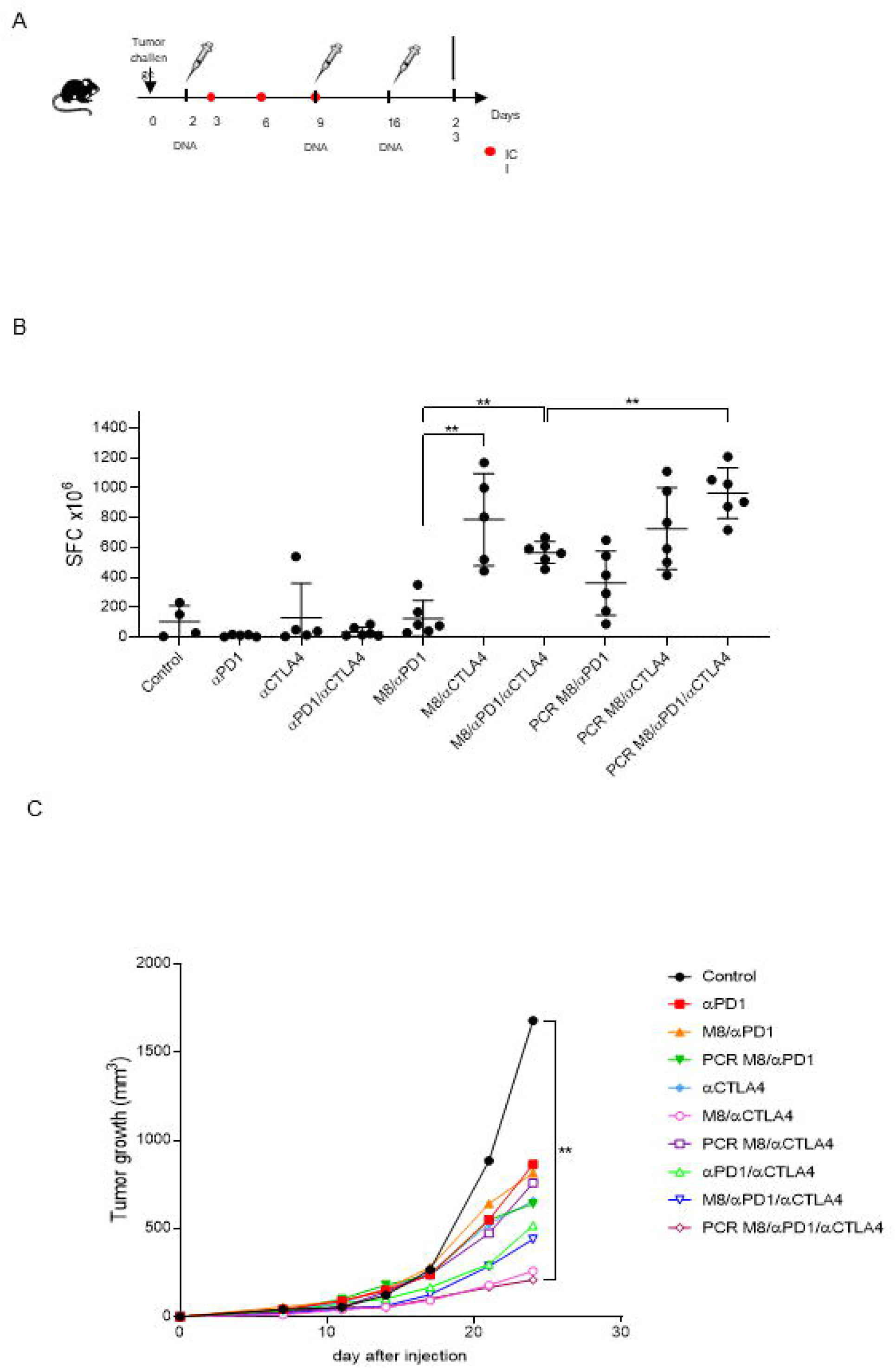
In vivo antitumoral effect of DNA amplicons. **(A)** Schematic representation of experimental setup in a neoantigen cancer murine model. After tumor challenge, C57Bl/6 mice received a combination of DNA (plasmid or amplicon, administered at days 2,9 and 16) and ICIs (αCTLA-4 or αPD1, administered at days 3,6 and 9). **(B)** Neoantigen-specific T cell response was assessed by IFNγ ELISpot performed on splenocytes collected at day 23 and stimulated with neoantigen peptide pools. Significance was determined using Mann-Whitney test, *p<0,05 **p< 0.01. **(C)** Panel depicts tumor growth curve up to sacrifice at day 23.

## Discussion

The concept of DNA-based vaccines is an appealing immunotherapeutic approach in many diverse fields, potentially translatable to the clinics, thanks to several beneficial features. In fact, DNA vaccines are engineered for maximal gene expression and immunogenicity and they allow for fast and scalable manufacturing as well as long-term stability at room temperature. Moreover, DNA vaccines do not require complex formulations such as those based on nanoparticles (necessary for peptide- or RNA-based vaccines) and have shown a highly satisfactory safety profile, without the potential risk of integration or pathogenicity.^24^ Importantly, they can be quickly designed from new genetic viral sequences and this technical advantage may be essential not only to respond to the emergency outbreak of an epidemic^25,26^ but also to generate personalized vaccines for treating tumors.^27^ In order to gain an efficient DNA uptake, new delivery technologies may accelerate the rapidly growing sector of genetic cancer vaccines. Among others, electroporation significantly increases the initial uptake of DNA by local cells and for this reason it has been widely used in different biomedical applications.^11^

Nonetheless, one huge hurdle of DNA vaccines manufacturing is their production as plasmids, grown in genetically modified bacteria containing, besides the gene of interest, a bacterial origin of replication and a selective gene, normally encoding for an antibiotic resistance, in order to maintain the persistence of the plasmid in the bacterium. Given the excessive and often inappropriate use of antibiotics, both in human and veterinary medicine, as well as in animal husbandry and agriculture, the last decades have witnessed the spread of these compounds in the environment on a global scale and consequently the onset of multidrug-resistant bacterial pathogens with evident and worrying effects on public health.^28,29^ In response to recent concerns about the antibiotic resistance spread associated with conventional DNA vaccines, alternative technologies have been proposed and new DNA vaccine technologies have recently emerged in which the DNA is manufactured in a cell-free process that avoids bacterial fermentation and yields a vaccine that is structurally linear, such as minicircles based on closed linear DNA.^30^ These new bacteria-free manufacturing platforms have already proven to be stable and immunogenic in preclinical models.^31,32^

Here we describe, for the first time in literature, the use of an innovative cell-free manufacturing platform as cancer immunotherapy in preclinical tumor model. In this study, enzymatically produced amplicon expression vectors have been specifically used as a DNA cancer vaccine to express tumor associated or mutated antigens able to elicit a specific immune response. As a proof of concept, a “universal” cancer DNA vaccine was produced comprising an amplicon expression vector encoding an optimized version of telomerase reverse transcriptase (TERT). High levels of TERT are detected in over 85% of cancers. TERT expression correlates with telomerase activity, which is required for tumor survival and unlimited proliferative capacity of cancer cells. In addition, we explore a personalized approach, i.e. DNA vaccines targeting tumor neoantigens. Neoantigens arise from somatic mutations that differ from wild-type antigens and are specific to each individual subject, or a sub-population of subjects, which provide tumor specific antigen targets. Neoantigens are found only in tumor cells, and thus are not subject to self-tolerance, have a decreased risk of generated autoimmunity and on-target off-tumor effects. Tumor neoantigens may be identified by differential sequencing of a subject’s tumor versus wild-type samples, using exome/genome sequences and RNAseq analysis, and the assistance of artificial intelligence, machine learning and predictive algorithms. Through this method, a DNA cancer vaccine comprising one or more amplicon expression vectors encoding the identified neoantigens can be designed and produced that will elicit an antigen-specific immune response to said neoantigens, resulting in a cancer vaccine with limited off-tumor side effects and high efficacy. Such “personalized” neoantigen DNA cancer vaccines are well suited for amplicon expression vectors and their method of rapid manufacture. With modern next generation sequencing (e.g. Illumina NovaSeq 6000), efficient bioinformatics platforms and rapid high-fidelity synthetic DNA synthesis, is it possible to create a therapeutic dose of personalized neoantigen encoding amplicon expression vectors in under 48 hours.

In a first phase, we looked at levels and length of gene expression, using luciferase as biomarker. As described in Fig. 2 and 3, the PCR-produced amplicons are highly stable *in vivo* over time, keeping a constant luciferase expression level up to 48 hours, then their expression capacity starts declining, although expression may be still detectable one week post injection. In addition, through the incorporation of chemical and/or peptide modifications, PCR-produced amplicons can be optimized for high-level expression within target cells leading to an enhanced antigen-specific immune response. In this perspective, it has been shown here that vaccination of mice with DNA amplicons induces similar levels of antigen-specific T cell response as compared with plasmid DNA. Particularly, as previously observed,^33^ a strong polyfunctional cell-mediated immune response was elicited by amplicons (Fig.4), as well as plasmids, thus validating this innovative technology as a valid alternative to DNA plasmids in the context of genetic vaccines.

Moreover, besides showing strong *in vivo* stability and immunogenicity, DNA amplicons have been shown to induce a robust antitumoral immune response specific for neoantigens expressed by a colon cancer model. It has been already shown elsewhere that neoantigen-based DNA vaccine approach is a potential alternative to currently established cancer immunotherapies, especially when administered as cotreatment with immune checkpoint inhibitors.^5^ By demonstrating that DNA amplicons encoding for neoantigens are equally effective as plasmids in antitumoral cotreatment with ICIs (Fig.5), we here validate the use of amplicon expression vectors as DNA cancer vaccines, as a more cost- and time-effective alternative to conventional plasmids, given their enhanced ability to rapidly manufacture tumor-specific cancer vaccines able to elicit antigen-specific immune responses with increased efficacy and reduced on-target off-tumor effects. Taken together, these results provide the first *in vivo* preclinical demonstration of linear DNA amplicons antitumoral efficacy thus encouraging further studies for clinical applications.

## Acknowledgments

Research activities are supported in part by the Italian Ministry of Economic Development through grant F/190180/01/X44 and Campania Regional grant B61G18000470007.

## Author contributions

A.C., E.S., L.L., M.C., E.P. performed experiments; A.C., F.P and L.A. analyzed and interpreted data; A.C. prepared the figures; L.A. provided funding, conceptual advice and edited the manuscript; A.C. coordinated the study and wrote the paper; B.V., Y.S. and B.L. produced the amplicons; J.H. and C.S. provided conceptual advice, coordinated the study and edited the manuscript.

## Data and materials availability

All data and materials are available in the main text.

